# BCL11A interacts with SOX2 to control the expression of epigenetic regulators in lung squamous cell carcinoma

**DOI:** 10.1101/223776

**Authors:** Kyren A. Lazarus, Fazal Hadi, Elisabetta Zambon, Karsten Bach, Maria-Francesca Santolla, Julie K. Watson, Lucia L. Correia, Madhumita Das, Rosemary Ugur, Sara Pensa, Lukas Becker, Lia S. Campos, Graham Ladds, Pentao Liu, Gerard Evan, Frank McCaughan, John Le Quesne, Joo-Hyeon Lee, Dinis Calado, Walid T. Khaled

**Affiliations:** Department of Pharmacology, University of Cambridge, Cambridge, UK; Department of Biochemistry, University of Cambridge, Cambridge, UK; Department of Pharmacy, Health and Nutritional Sciences, University of Calabria, Rende, Italy; Wellcome Trust Sanger Institute, Cambridge, UK; The Francis Crick Institute, London, UK; WT-MRC Stem Cell Institute, University of Cambridge, UK; Cambridge Cancer Centre, Cambridge, UK; MRC Toxicology Unit, Lancaster Road, Leicester; Cancer Research Centre, University of Leicester; University Hospitals Leicester NHS trust

## Abstract

Patients diagnosed with lung squamous cell carcinoma (LUSC) have limited targeted therapeutic options. We report here the identification and characterisation of the transcriptional regulator, *BCL11A*, as a LUSC oncogene. Analysis of cancer genomics datasets revealed *BCL11A* to be upregulated in LUSC but not lung adenocarcinoma (LUAD). We demonstrated that knockdown of *BCL11A* in LUSC cell lines abolished xenograft tumour growth and its overexpression *in vivo* led to lung airway hyperplasia and the development of reserve cell hyperplastic lesions. In addition, deletion of *Bcl11a* in the tracheal basal cells abolished the development of tracheosphere organoids while its overexpression led to solid tracheospheres with a squamous phenotype. At the molecular level we found *BCL11A* to be a target of SOX2 and we show that it is required for the oncogenic role of SOX2 in LUSC. Furthermore, we showed that *BCL11A* and SOX2 interact at the protein level and that together they co-regulated the expression of several transcription factors. We demonstrate that pharmacological inhibition of SETD8, a gene co-regulated by BCL11A and SOX2, alone or in combination with cisplatin treatment, shows significant selectivity to LUSC in comparison to LUAD cells. Collectively, these results indicate that the disruption of the BCL11A-SOX2 transcriptional program provides a future framework for the development of targeted therapeutic intervention for LUSC patients.

## Main

Lung cancer accounts for the highest rate of cancer related diagnosis and mortality worldwide^1^. Broadly, there are two major types of lung cancer; small cell lung cancer (SCLC) accounting for 15% of the cases and non-small cell lung cancer (NSCLC) accounting for 85% of cases^1^. NSCLC patients have a poor outcome in the clinic with only 15% of patients surviving five years or more^2^. At present NSCLC is defined histo-pathologically in the clinic into four broad categories: lung adenocarcinoma (LUAD), lung squamous cell carcinoma (LUSC), large cell carcinoma and undifferentiated non-small cell lung cancer. LUAD and LUSC are the most common types of NSCLC (90% of cases). LUSC accounts for more than 400,000 deaths worldwide each year^3^ and unlike LUAD there are limited targeted therapies available for LUSC. Platinum-based chemotherapy remains the first-line treatment for LUSC and although the recent FDA approval of Necitumumab in combination with platinum-based chemotherapy for metastatic LUSC has shown positive signs, a great deal of work still needs to be done in this field^4^.

At the cellular level, LUAD tends to originate from the secretory epithelial cells in the lung while LUSC usually originates from basal cells in the main and central airways^2^. At the molecular level LUAD is known to harbour mutations in epidermal growth factor receptor (EGFR), V-Ki-Ras2 Kirsten Rat Sarcoma Viral Oncogene Homolog (KRAS) and Anaplastic Lymphoma Receptor Tyrosine Kinase (ALK), which are also well modelled and studied both *in vitro* and *in vivo* (for review see ^5,6^). On the other hand LUSC is less studied but it is known that amplifications of SRY (Sex Determining Region Y)-Box 2 (SOX2) tends to be present in 70-80% of patients^7–10^.

Recently, a detailed picture of the molecular differences between LUAD and LUSC has been made available through ‘The Cancer Genome Atlas’ (TCGA)^11,12^. To identify key drivers responsible for the differences between LUAD and LUSC we reanalysed the gene expression data from TCGA and focused on all 1500 transcriptional regulators in the genome. As reported previously *SOX2* was the most amplified gene in LUSC and its expression level was also significantly higher in LUSC vs. LUAD (Figure 1a and Supplementary Fig. 1a). The second most amplified locus in LUSC patients revealed by TCGA analysis contains the transcription factors *BCL11A* and *REL^11,12^. BCL11A* has been shown to be an oncogene in B-cell lymphoma and triple negative breast cancer^13–16^.

**Figure 1.**
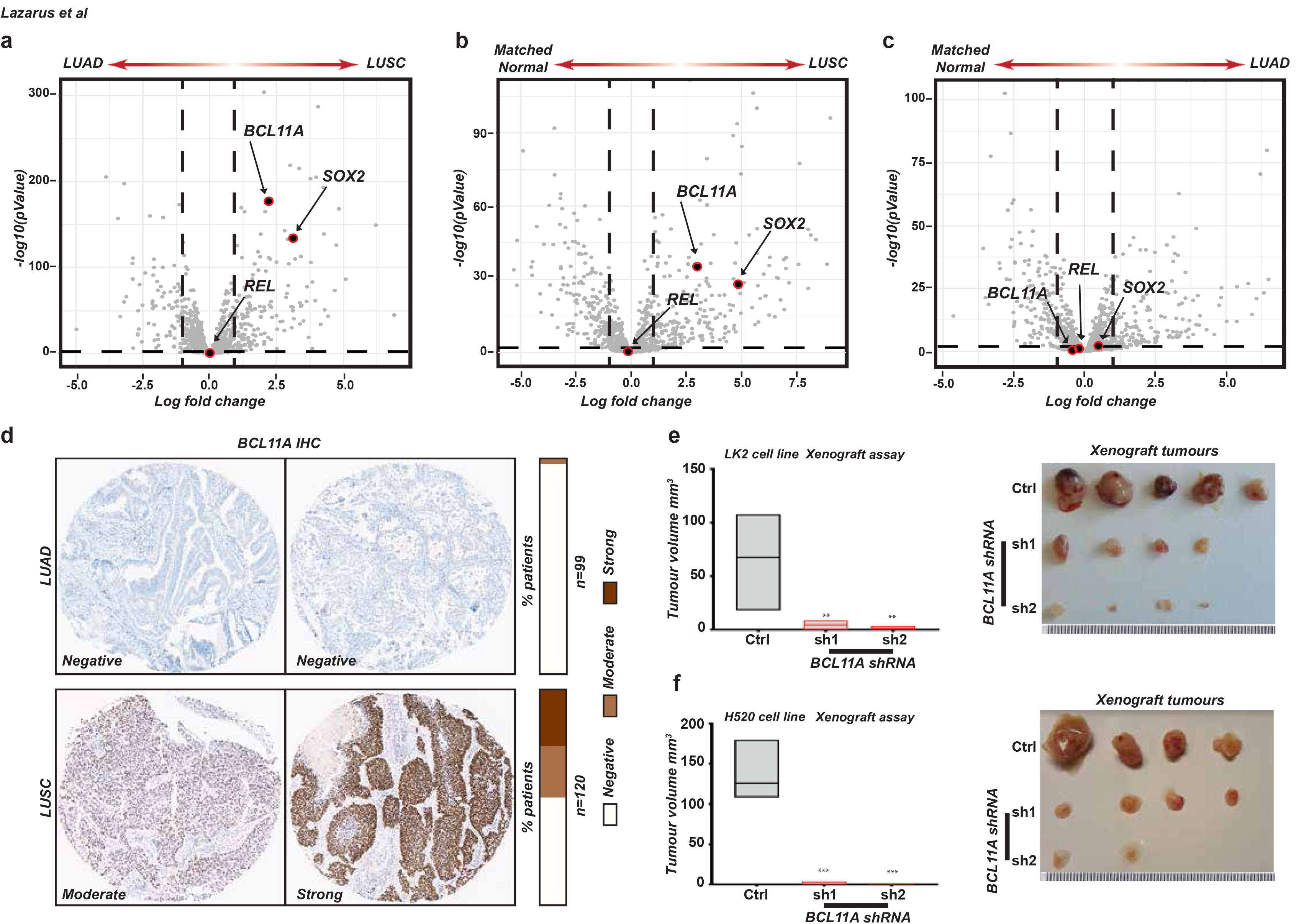
*BCL11A* is a lung squamous cell carcinoma (LUSC) oncogene. **(a)** Volcano plots of The Cancer Genome Atlas (TCGA) RNAseq data^11,12^ indicating that *BCL11A* and *SOX2* are higher expressed in LUSC compared to lung adenocarcinoma (LUAD). The plot shows that *REL* is not differentially expressed in LUSC vs LUAD patients. The x-axis represents log2 expression fold-change (FC) in LUSC patients vs LUAD patients and the y-axis represents –log10(pValue). The vertical dashed lines represent FC = 1.0 and the horizontal dashed line represents p value = 0.01. **(b)** Volcano plots indicating that *BCL11A* and *SOX2* are differentially expressed in LUSC patients vs matched normal samples. The plot indicates that *REL* is not differentially expressed in LUSC vs matched normal samples **(c)** Volcano plots indicating that neither BCL11A, SOX2 nor *REL* are differentially expressed in LUAD patients vs matched normal. **(d)** Images and scoring of BCL11A IHC staining on LUAD and LUSC tumours (see Methods for scoring). Graph depicting reduction in tumour size observed when shRNA1 or shRNA2 transfected LK2 **(e)** and H520 **(f)** cells are injected subcutaneously into mice compared to control. Five mice per cell line were monitored for 25 days after which tumours were removed and measured. On the right are images showing actual tumours measured. Data presented as mean ± s.d. One way ANOVA with post Dunnett test performed, * indicates p<0.05 and ** p<0.005 and *** indicates p<0.001.

We found that *BCL11A* expression levels were also significantly higher in LUSC vs. LUAD (Figure 1a and Supplementary Figure 1a). Moreover, the expression of both *BCL11A* and *SOX2* was significantly higher in LUSC but not LUAD tumour samples compared to patient matched normal samples (Figure 1b-c and Supplementary Figure 1b-c) supporting a driver role for these transcription factors in LUSC pathology. In contrast, *REL* expression was not different between LUSC and LUAD (Figure 1a-c and Supplementary Figure 1a-c) suggesting that *BCL11A* amplification is a key driving event in LUSC. This observation is supported by the recent report from the TRACERx (TRAcking Cancer Evolution through therapy (Rx)) study demonstrating the amplification of *BCL11A* as an early event in LUSC^17^. Furthermore, BCL11A IHC staining on LUAD (n=99) and LUSC (n=120) TMAs revealed little or no staining in 99% of LUAD patients. In contrast, 25% of LUSC patients had moderate staining and 24% of LUSC patients had strong staining, which is in agreement with previous IHC staining of NSCLC tumours^18^. This confirms our transcriptomic analyses indicating the specificity of BCL11A in LUSC patients.

To determine if high levels of *BCL11A* expression are oncogenic in LUSC, we performed shRNA mediated knockdown (KD) of *BCL11A* using two independent shRNAs in two LUSC cell lines, LK2 and H520 (Supplementary Fig. 2a and b). We first tested the clonogenic capacity of control or *BCL11A-KD* cells by seeding them into matrigel for 3D colony formation assays. We found that *BCL11A-KD* cells had a significant reduction in colony formation capacity (Supplementary Fig. 2c and d). We then injected control or BCL11A-KD cells into immune compromised mice to test for their tumour formation capacity and found a significant reduction in tumour burden from *BCL11A-KD* cells compared to control cells (Fig 1e and f). In addition, we found the squamous markers *KRT5* and *TP63* levels were significantly reduced in *BCL11A-KD* cells (Supplementary Figure 2e-h) suggesting an integral role for *BCL11A* in driving the LUSC phenotype. Moreover, to test if the role of BCL11A is context dependant we knocked down *BCL11A* in an LUAD cell line H1792 and found no change in 3D colony growth (Supplementary Figure 2i-j). Finally, at the molecular level we found no change in the expression of *SOX2* in *BCL11A-KD* cells (Supplementary Figure 2k, l).

To explore the role of BCL11A in lung biology, we utilised a novel Cre-inducible mouse model that allows for the overexpression of *BCL11A*. Essentially, *BCL11A* was inserted into the *Rosa26* locus with a LoxP-Stop-LoxP (LSL) cassette upstream, under the control of a GAGG promoter, thus preventing the expression of *BCL11A* unless the *LSL* is excised by Cre recombinase. To test the effect of *BCL11A* overexpression on lung morphology, we allowed the BCL11A-overexpression (*BCL11A*^ovx^) mice to inhale Adenovirus-Cre (Figure 2a). Eight months after infection, we analysed the lungs and found an increase in airway hyperplasia (Figure 2b-c) accompanied by aggregates of small hyperchromatic cells with irregular nuclei that represent reserve cell hyperplasia (Figure 2b, arrows) which are precursors to squamous metaplasia^19^. IHC analysis of the lungs from *BCL11A^ovx^* also indicated an increase in positivity for the proliferative marker Ki-67 (Supplementary Figure 3a) and Sox2 indicating a transition to squamous differentiation (Supplementary Figure 3b). However, we found little difference in Cc10, Krt5 and Trp63 staining at this stage (Supplementary Figure 3a and b).

**Figure 2.**
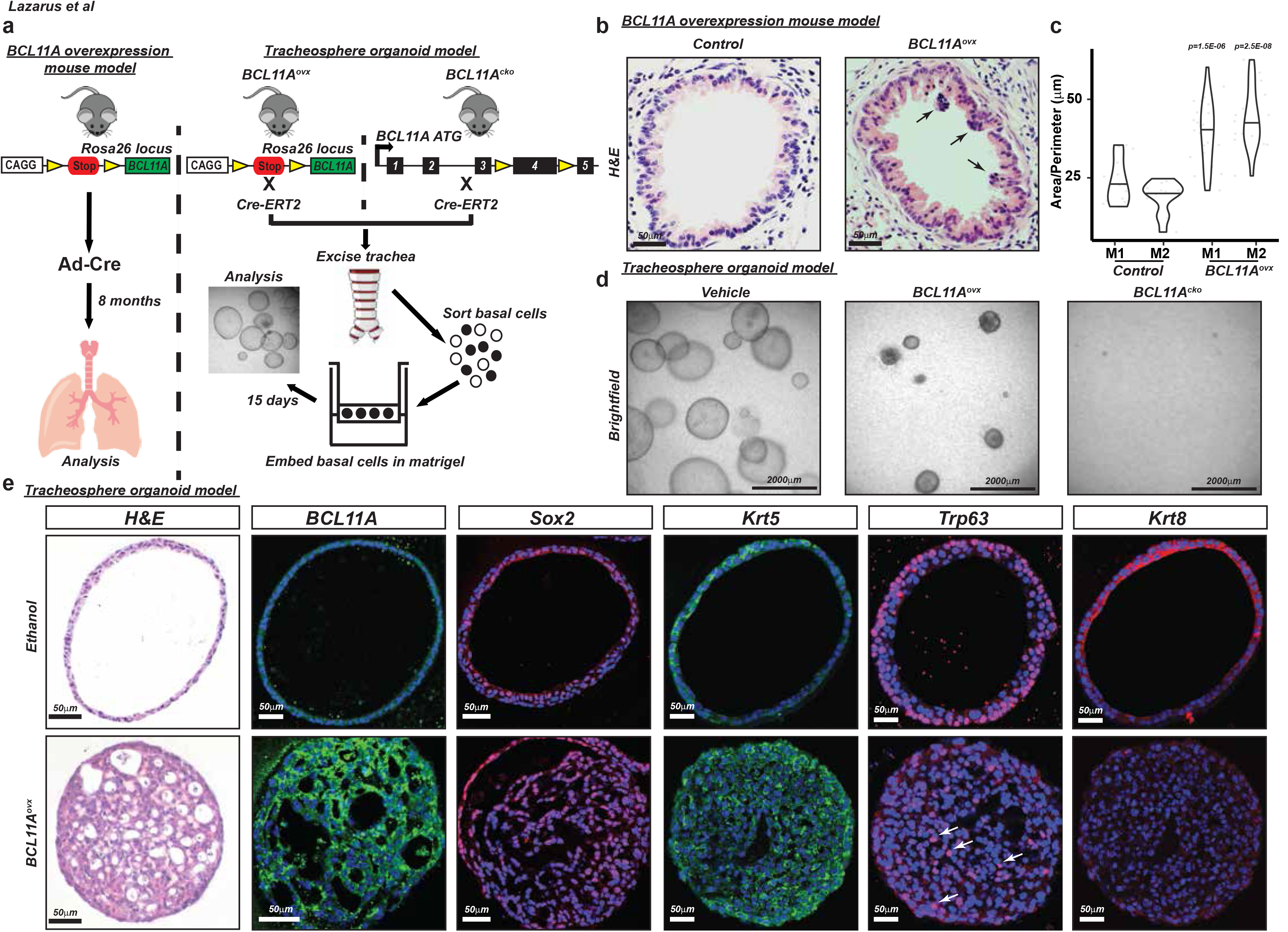
*BCL11A* overexpression leads to thickening of the airways and abnormal organoid formation. **(a)** Schematic representing strategy to explore the role of *BCL11A in vivo* and *ex vivo*. Left Panel: Adenovirus-Cre was nasally administered to *BCL11A^ovx^* mice and the lungs were analysed after eight months. Right panel: for the tracheosphere organoid model, basal cells from the trachea of either *BCL11A^ovx^* or *BCL11A^cko^* mice were FACS sorted, embedded in matrigel and analysed after 15 days. Three independent mice were used for each experiment. **(b)** Images of airways from control and *BCL11A^ovx^*. Arrows indicate small hyperchromatic cells with irregular nuclei. **(c)** Quantificaiton of airway epithelial layer hyperplasia from two control and *BCL11A^ovx^* mice. **(d)** Bright field images of organoids from *Bcl11a^cko^* and *BCL11A^ovx^* mice treated with vehicle or tamoxifen. **(e)** Sectioned organoids from *BCL11A^ovx^* mice stained with haematoxylin and eosin, BCL11A, Sox2, Krt5, Trp63 and Krt8. Scale bar indicates 50μm.

To further investigate the role of *BCL11A* in LUSC, we employed a recently described 3D organoid culture system for basal cells (BCs) from the trachea, as they have been suggested to be the cell of origin for LUSC^20^. BCs from human and mouse lungs have higher expression of *BCL11A* when compared to the other epithelial compartments (Supplementary Figure 4a and b)^21^. Therefore, we crossed the *BCL11A^ovx^* mice to *R26-CreERT2* mice, which allowed us to induce Cre recombination by the administration of tamoxifen. In addition, we also used the previously reported *Bcl11a* conditional knockout mouse under the control of *R26-CreERT2* (called *Bcl11a^CKO^* from this point) to elucidate the importance of BCL11A in organoid formation (Figure 2a).

We found that in contrast to the hollow organoids normally formed by wild-type BCs, BCs from *BCL11A^ovx^* mice formed solid organoid structures with no hollow lumen suggesting hyper-proliferation and loss of organisation (Figure 2d). However, BCs from *Bcl11a^cko^* mice failed to form any organoid structures suggesting that *Bcl11a* is necessary for stem cell potential in organoid formation (Figure 2d). Quantitative analysis revealed a significant decrease in organoid numbers from *Bcl11a^cko^* BCs but no significant difference in *BCL11A^ovx^* BCs (Supplementary Figure 4c-e).

H&E staining confirmed the hollowness of the organoids from the control mice which was in stark contrast to the filled organoids from the *BCL11A^ovx^* mice (Figure 2e and Supplementary Figure 5). IF staining revealed that the *BCL11A^ovx^* organoids were also positive for Ki-67 (Supplementary Figure 5), Sox2, Krt5, Trp63 and negative for the luminal marker Krt8 (Figure 2e and Supplementary Figure 6) indicating that BCL11A maintains a squamous phenotype.

Given the importance of SOX2 in driving LUSC^8,9^ we next investigated if *BCL11A* is regulated by SOX2. To achieve this, we knocked down *SOX2* (*SOX2-KD*) in two LUSC cell lines using two independent shRNAs (Figure 3a, b and Supplementary Figure 7a, b). Similar to *BCL11A* we found that *SOX2-KD* cells had a significantly reduced colony and tumour formation abilities (Supplementary Figure 7c-f). At the molecular level we found a significant reduction in the expression levels of *BCL11A* and similar to *BCL11A-KD*, reduction in squamous markers *KRT5* and *TP63* in the *SOX2-KD* cells (Figure 3c, d and Supplementary Figure 7g-j).

**Figure 3.**
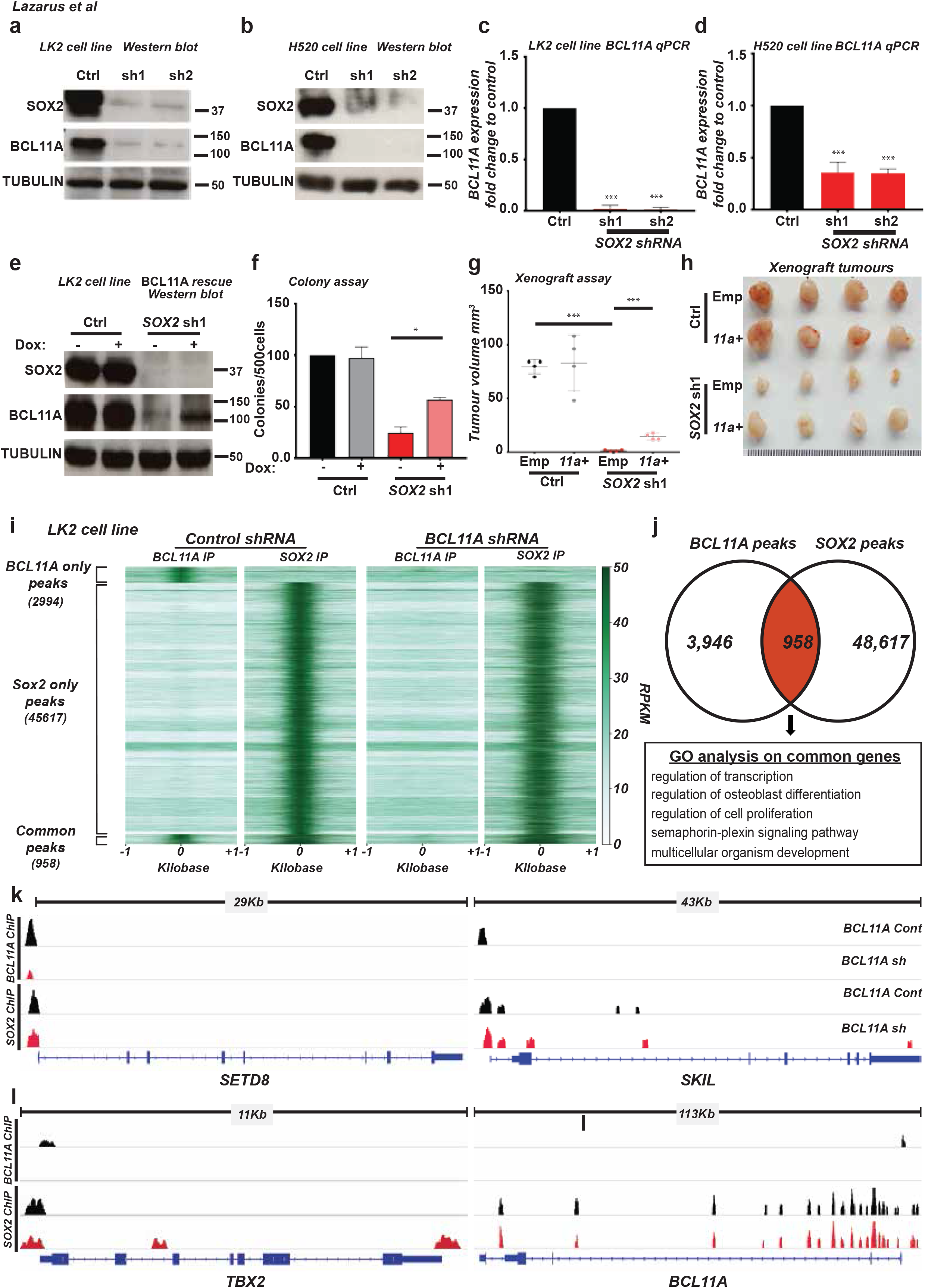
BCL11A and SOX2 occupy independent and common loci in the genome of LUSC cells. **(a and b)** Western blot showing SOX2 and BCL11A expression in *SOX2-KD* in LK2 **(a)** and H520 **(b)** cells transfected with control (scramble), shRNA1 or shRNA2 vectors. **(c and d)** *BCL11A* expression in SOX2-KD *LK2* **(c)** and *H520* **(d)** cells. **(e)** Western blot showing BCL11A rescue in SOX2-KD cells. Doxycycline (Dox) inducible *BCL11A* overexpression vector was transfected into control and *SOX2* shRNA1 LK2 cells and Dox treatment was performed for 48 hours. **(f)** Graph depicting 3D matrigel assay in control, *SOX2-KD* and *BCL11A* rescue cells indicating a partial rescue in *SOX2-KD, BCL11A* overexpressing cells. **(g)** Graph indicating partial rescue of tumour size from *BCL11A* overexpressing *SOX2-KD* cells injected subcutaneously. **(h)** Images of actual tumours measured. Four mice per cell line were monitored for 15 days after which tumours were removed and measured. Data presented as mean ± s.d. (n=4). One way Anova test performed, * indicates p<0.05 and ** p<0.005 and *** indicates p<0.001. **(i)** LK2 cell line either transfected with control or shRNA1 vectors were used for BCL11A and SOX2 ChIP-Seq. Heatmaps showing BCL11A only, SOX2 only or common peaks in BCL11A or SOX2 IP in control and *BCL11A*-KD cells. **(j)** Venn diagram indicating the overlap of BCL11A and SOX2 target genes in LK2 cells. BCL11A target genes were derived by comparing BCL11A IP in control vs BCL11A-KD cells. Any peaks lost in BCL11A-KD cells were considered as a ‘true peak’ and included as a bona fide BCL11A target gene. SOX2 target genes were derived from SOX2 IP in LK2 control cells. Image below show top five biological GO terms from GO analysis performed using DAVID. **(k)** IGV genome browser views for *SETD8, SKIL, TBX2* and *BCL11A*.

To investigate if BCL11A is required for SOX2 mediated oncogenesis we introduced a doxycycline inducible *BCL11A* overexpression vector into *SOX2-KD* cell lines and found that *BCL11A* overexpression partially rescues the colony and tumour formation abilities of *SOX2-KD* cells (Figure 3e-h, f and Supplementary Figure 8a-c). These results suggest that BCL11A is partially responsible for some of SOX2’s mediated transcriptional changes in LUSC cells.

To understand how BCL11A and SOX2 contribute to the LUSC transcriptional programme we performed BCL11A and SOX2 ChIP-Seq analysis on LK2 cells in the presence or absence of BCL11A (*BCL11A-KD*) (Figure 3i). We identified 49,575 peaks for SOX2 and 4,904 peaks for BCL11A (Figure 3j and Supplementary Tables 1 and 2). Out of the 4,904 BCL11A peaks identified, 3,941 were not present in *BCL11A-KD* cells validating the true nature of BCL11A binding at these regions of the genome.

We then compared the BCL11A peaks and the SOX2 peaks and identified 958 peaks, which overlap suggesting co-regulation of these genes (Figure 3i and j). Subsequently, we identified the nearest genes to the common peaks (Supplementary Table 3) and performed Gene Ontology (GO) analysis to identify if these common peaks are enriched for specific biological process (Figure 3j). The top hits from the GO analysis revealed enrichment for transcriptional and epigenetic regulators including *SETD8, SKIL*, and *TBX2* (Figure 3k). We also found that SOX2 binds the *BCL11A* locus at multiple sites suggesting a strong direct regulation at the transcriptional level further supporting the data in Figure 3c and d (Figure 3k). BCL11A and SOX2 peaks on *SETD8^22,23^, SKIL^24^, TBX2*^25^ and *BCL11A* were validated and confirmed by ChIP-qPCR (Supplementary Figure 9). The overlap of the ChIP-Seq peaks suggests a direct interaction between BCL11A and SOX2 proteins, which was confirmed in co-immunoprecipitation experiments on LK2 and H520 (Supplementary Figure 10a, b).

We then investigated the expression of *SETD8, SKIL* and *TBX2* in *BCL11A-KD* or *SOX2-KD* cells. Interestingly, we found a modest reduction in the expression of these genes in *BCL11A-KD*, which was more pronounced in *SOX2-KD* cells (Figure 4a-d; and Supplementary Figure 10c-j). This indicates that SOX2 has a major role in the regulation of *BCL11A*, which together with SOX2 drives the expression of key epigenetic regulators such as *SETD8, SKIL* and *TBX2*. This results suggest that disrupting the BCL11A-SOX2 transcriptional program might selectively target LUSC cells. To test this hypothesis we focused on SETD8 for its biology and the availability of multiple inhibitors. SETD8 is a member of the SET domain containing family and is known to catalyse the monomethylation of histone H4 Lys20^22,23^ which is involved in recruiting signalling proteins or chromatin modifications^26^. In addition, SETD8 has been shown to play a role in maintaining skin differentiation^27^ and is dysregulated in multiple cancer types^28-30^.

**Figure 4.**
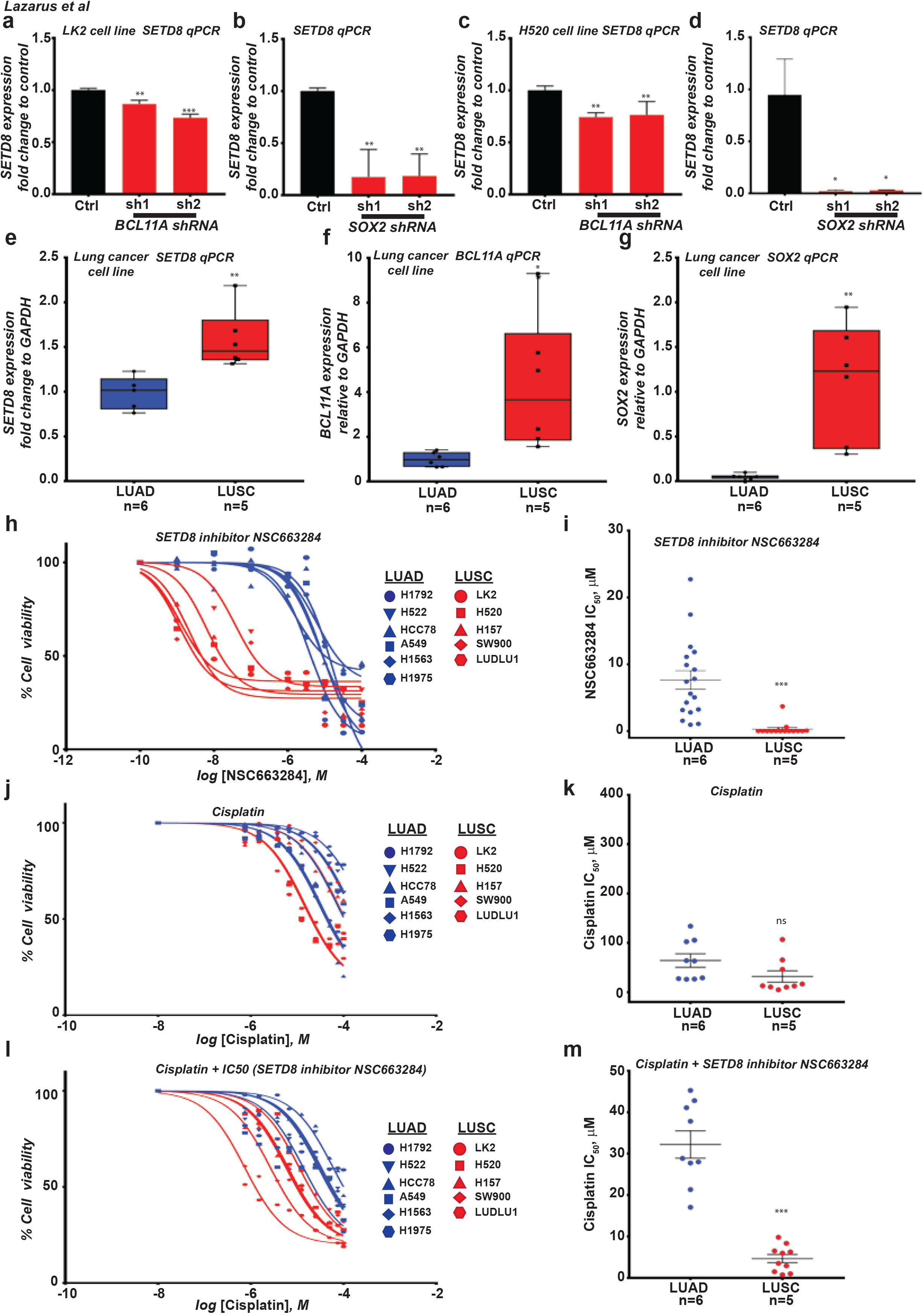
SETD8 inhibition preferentially sensitises LUSC cell lines to chemotherapy. **(a-d)** Graph depicting *SETD8* gene expression in LK2, H520 *BCL11A-KD* and LK2, H520 *SOX2 KD* cells. Data presented as mean ± s.d. (n=3). One way ANOVA with post Dunnett test performed, * indicates p<0.05 and ** p<0.005 and *** indicates p<0.001. **(e)** *SETD8* gene expression in NSCLC cell lines. **(f)** *BCL11A* gene expression in NSCLC cell lines. **(g)** *SOX2* gene expression in NSCLC cell lines. **(h)** Dose-response curves were derived by treating NSCLC cell lines with SETD8 inhibitor NSC663284. 1000 cells were seeded and allowed to recover for 24 hours. The inhibitor was then added at increasing concentrations to LUSC (red) and LUAD (blue) cells and Cell Titre (see Methods) assay was performed after 72 hours. **(i)** IC_50_ values were derived from the dose-response assay indicating LUSC cells are significantly more responsive to SETD8 inhibition than LUAD cells. **(j)** Dose-response curves were derived by treating NSCLC cell lines with cisplatin as above. **(k)** IC_50_ values were derived from the dose-response assay indicating cisplatin effects LUSC and LUAD cells equally. **(l)** Dose-response curves derived from treating NSCLC cell lines with cisplatin and NSC663284 IC50 concentration for each cell line as above. **(m)** IC_50_ values were derived from the dose-response assay indicating SETD8 inhibition preferentially enhances cisplatin efficacy in LUSC cells. Data presented as mean ± s.d. (LUSC n=5 and LUAD n=6). Student’s t-test performed, * indicates p<0.05 and ** p<0.005 and *** indicates p<0.001.

We first assessed the expression of *SETD8* in multiple NSCLC cell lines and found that *SETD8* expression was significantly higher in LUSC cell lines (n=5) compared to LUAD cell lines (n=6) (Figure 4e) which was in correlation with the expression levels of *BCL11A* and *SOX2* (Figure 4f and g). Analysis of TCGA datasets revealed that SETD8 expression correlates with BCL11A and SOX2 expression in LUSC patients (Supplementary figure 11a and b). Next, we tested if LUSC lines are more sensitive to SETD8 inhibition. There are few SETD8 inhibitors but we focussed on one with proven cellular activity^31,32^, named NSC663284. NSC663284 has also been reported as an inhibitor of Cdc25^33–35^. We treated all the NSCLC cell lines (n=11) with a range of NSC663284 concentrations (full details of setup in materials and methods) for 72 hours and measured cell viability. Remarkably, we found that LUSC cells had significantly lower IC_50_ (average 0.30 μM) compared to LUAD cells (average 7.65 μM) (Figure 4h and i). To understand if SETD8 inhibition would add a clinical benefit to patients, we tested the effect of combining NSC663284 and cisplatin. First, we found that cisplatin treatment for 24hrs alone affected LUSC and LUAD in a similar way with both cell types demonstrating a similar IC_50_ (LUAD = 64.12 μM and LUSC = 31.67 μM) (Figure 4j and k). However, if we pre-treat NSCLC cell lines with NSC663284 for 48hrs and then combine cisplatin with NSC663284 for a further 24hrs we found that LUSC (IC_50_ = 4.66 μM) cells are more sensitive to cisplatin than LUAD cells (IC_50_ = 32.21 μM) (Figure 4l, m).

In summary, we have demonstrated in this study that BCL11A is a LUSC oncogene. We have shown that *BCL11A* is a key target of SOX2 and together they co-regulate the expression of epigenetic regulators in LUSC cells. Our data suggests that disrupting the BCL11A-SOX2 transcriptional program might selectively target LUSC cells. This was demonstrated by the sensitivity of LUSC cell lines to the inhibition of the BCL11A/SOX2 co-regulated gene *SETD8*. Collectively, these results indicate the key oncogenic role of *BCL11A* in LUSC and provide a future framework for the development of targeted therapeutic intervention for LUSC patients.

**Supplementary Figure 1.**
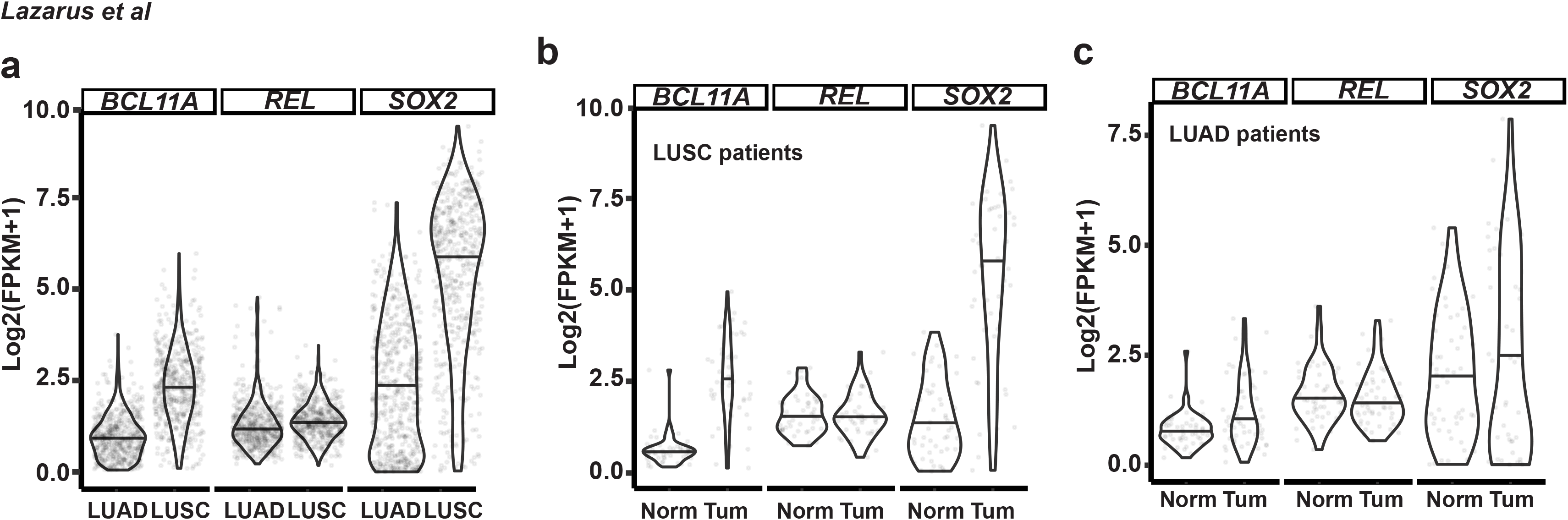
BCL11A, SOX2, and REL expression in Lung TCGA dataset. **(a)** Violin plot showing *BCL11A, SOX2* and *REL* expression in LUSC vs LUAD. FPKM, fragments per kilobase per million mapped reads. **(b)** Violin plot showing *BCL11A, SOX2* and *REL* expression in LUSC tumour vs normal matched patients. **(c)** Violin plot showing *BCL11A, SOX2* and *REL* expression in LUAD tumour vs normal matched patients.

**Supplementary Figure 2.**
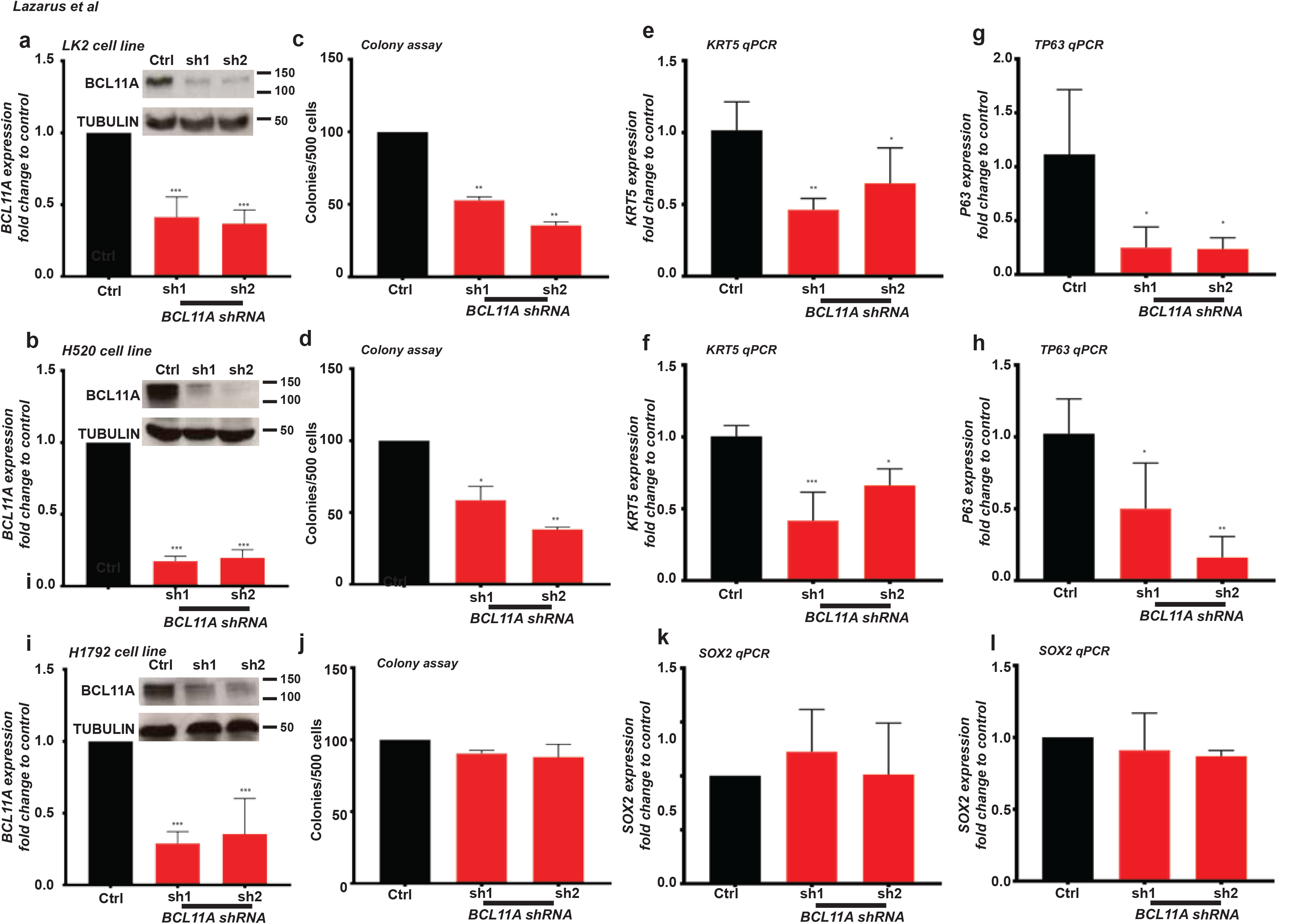
BCL11A KD reduces squamous markers in LUSC cells. **(a and b)** qPCR and western blot shows *BCL11A* reduction in LK2 **(a)** and H520 **(b)** cells transfected with shRNA1 and shRNA2 vectors. **(c and d)** Comparison of colony numbers in 3D matrigel assay from control, shRNA1 or shRNA2 in LK2 **(c)** and **(d)** H520 cells. Data presented as mean ± s. d. (n=3). **(e and f)** *KRT5* expression is reduced in LK2 **(e)** and H520 **(f)** *BCL11A-KD* cells. **(g and h)** *TP63* expression is reduced in LK2 **(g)** and H520 **(h)** *BCL11A-KD* cells. (i) qPCR and western blot shows BCL11A reduction in H1792 cells transfected with shRNA1 and shRNA2 vectors. **(j)** Comparison of colony numbers in 3D matrigel assay from control, shRNA1 or shRNA2 in H1792 cells. **(k and l)** *SOX2* expression is unchanged in LK2 **(k)** and H520 **(l)** *BCL11A-KD* cells. Data presented as mean ± s.d. One way ANOVA with post Dunnett test performed, * indicates p<0.05 and ** p<0.005 and *** indicates p<0.001.

**Supplementary Figure 3.**
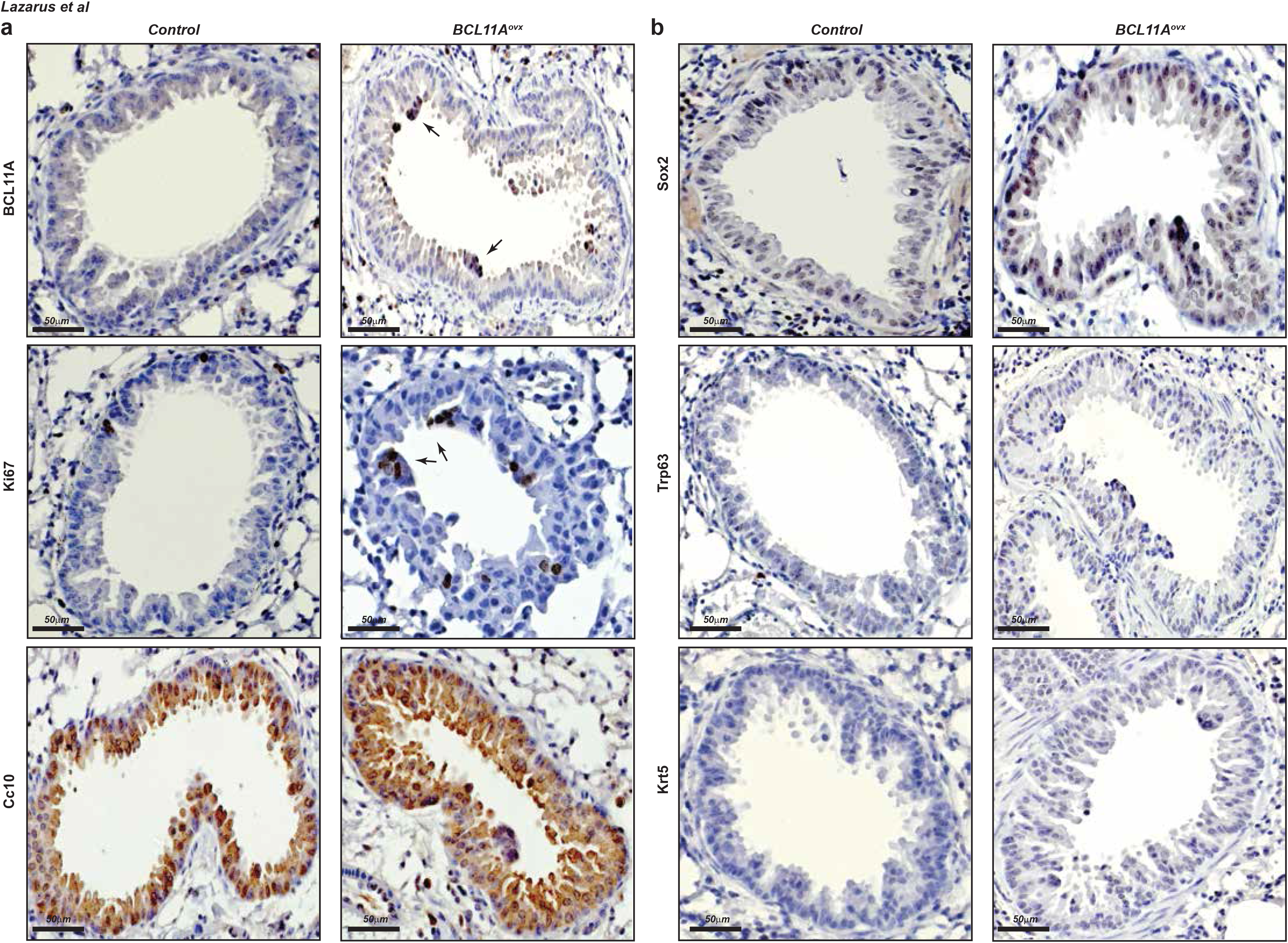
Airways from the *BCLL1A^ovx^* mice demonstrate proliferative preneoplastic lesions. **(a)** Immunohistochemistry for BCL11A, Ki67 and Cc10 expression in control vs BCL11A^ovx^ airways. BCL11A panel, arrows indicating positive staining especially in preneoplastic lesions. Ki67 panel, arrows indicating positive staining. **(b)** Sox2, Trp63 and Krt5 expression in control vs *BCL11A^ovx^* airways. Scale bar indicates 50 μm.

**Supplementary Figure 4.**
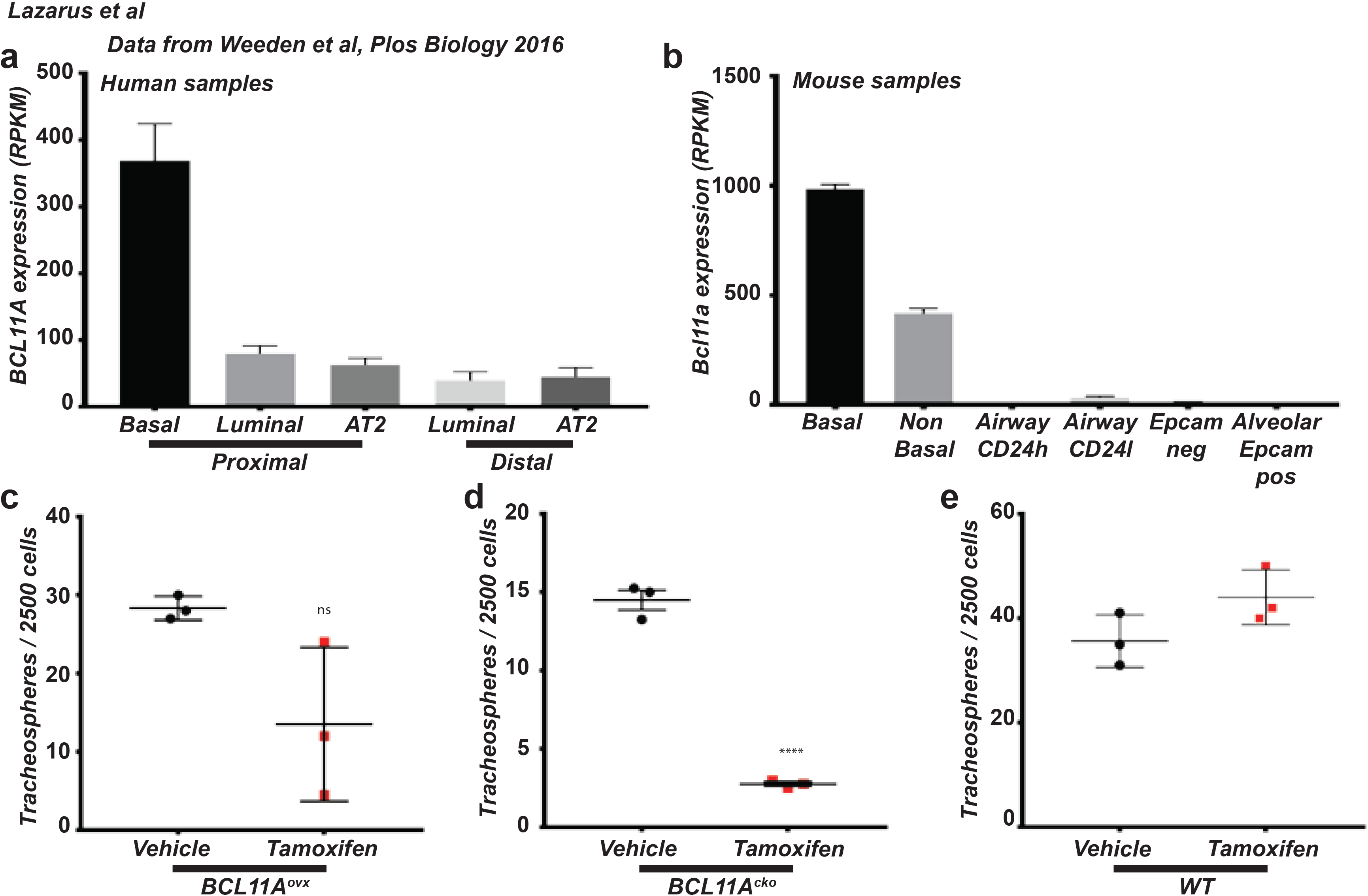
Organoids from *BCL11A^ovx^* basal cells exhibit increase in squamous markers. **(a)** RNA-Seq data analysed from Weeden et al ^21^ indicating that in human samples FACS fraction labelled as proximal BCs has approximately 400 fold increase in human *BCL11A* levels in comparison to other epithelial fractions. **(b)** RNA-Seq data from the same dataset indicating that mouse *Bcl11a* is approximately 500-1000 fold higher in basal fractions compared to other epithelial subtypes. **(c, d and e)** Quantification of organoids in vehicle vs tamoxifen treated **(c)** BCL11A^ovx^, **(d)** *BCL11A^cko^* and **(e)** WT organoids.

**Supplementary Figure 5.**
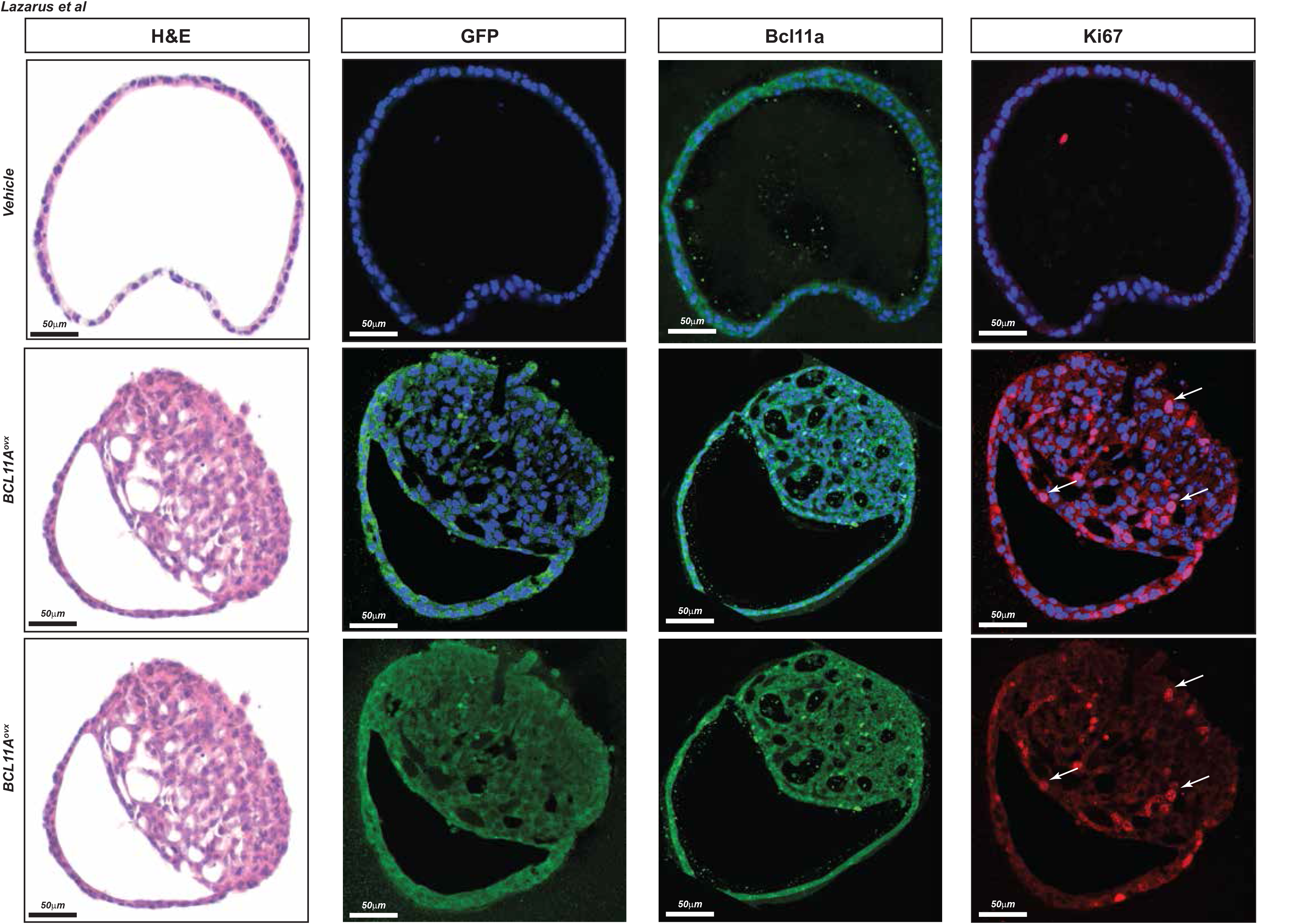
Organoids from the BCL11A^ovx^ basal cells display an abnormal proliferative phenotype. Vehicle and tamoxifen treated BCL11A^ovx^ organoids stained with H&E, GFP (which is also expressed if the LSL is efficiently excised), Bcl11a and Ki67. Nuclei stained by DAPI illustrated in blue. Arrows indicate positive staining. Scale bar indicates 50μm.

**Supplementary Figure 6.**
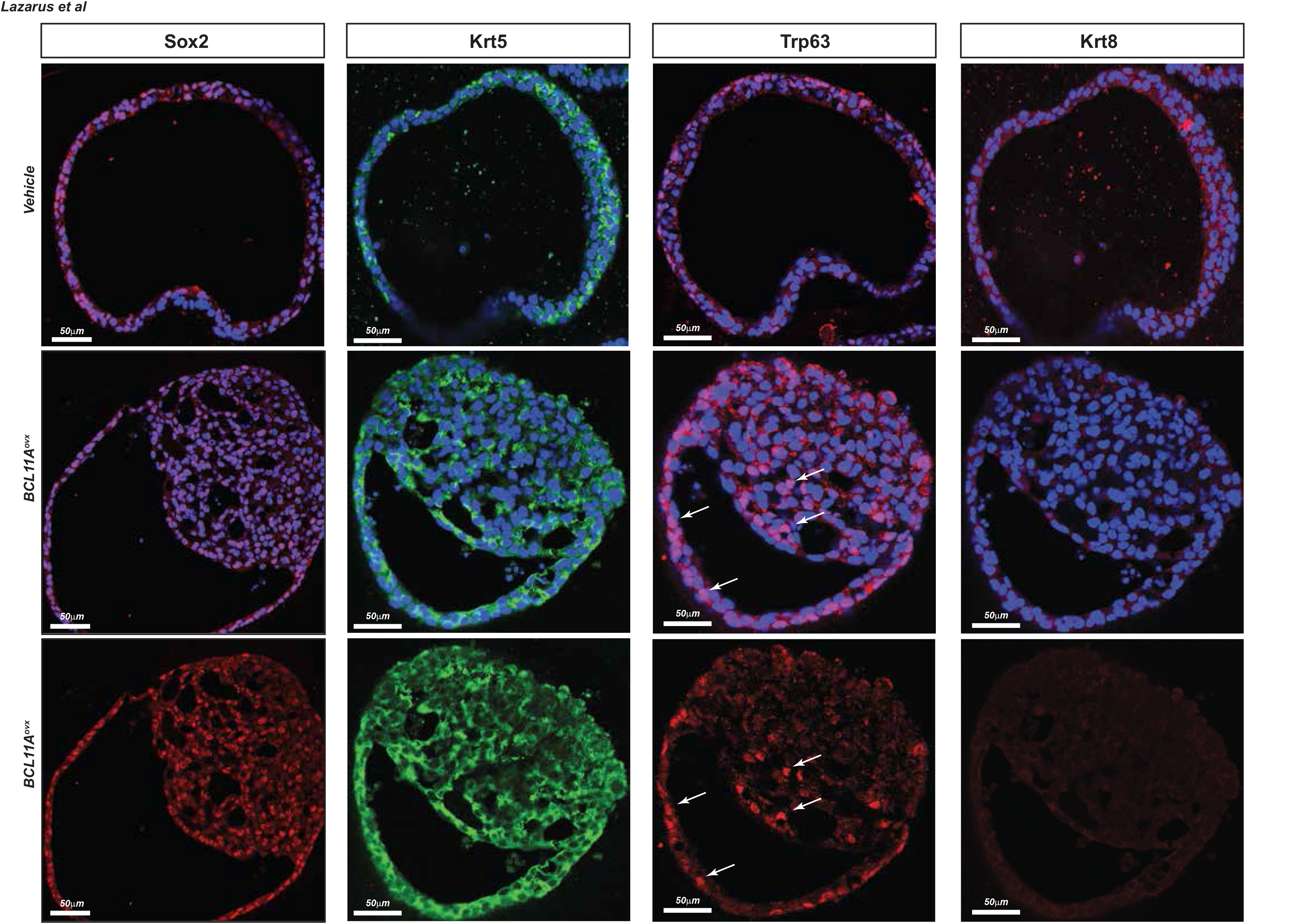
BCL11A overexpression inhibits organoid luminal differentiation. Vehicle and tamoxifen treated BCL11A^ovx^ organoids stained with Sox2, Krt5, Trp63 and Krt8. Nuclei stained by DAPI illustrated in blue. Arrows indicate positive staining. Scale bar indicates 50 μm.

**Supplementary Figure 7.**
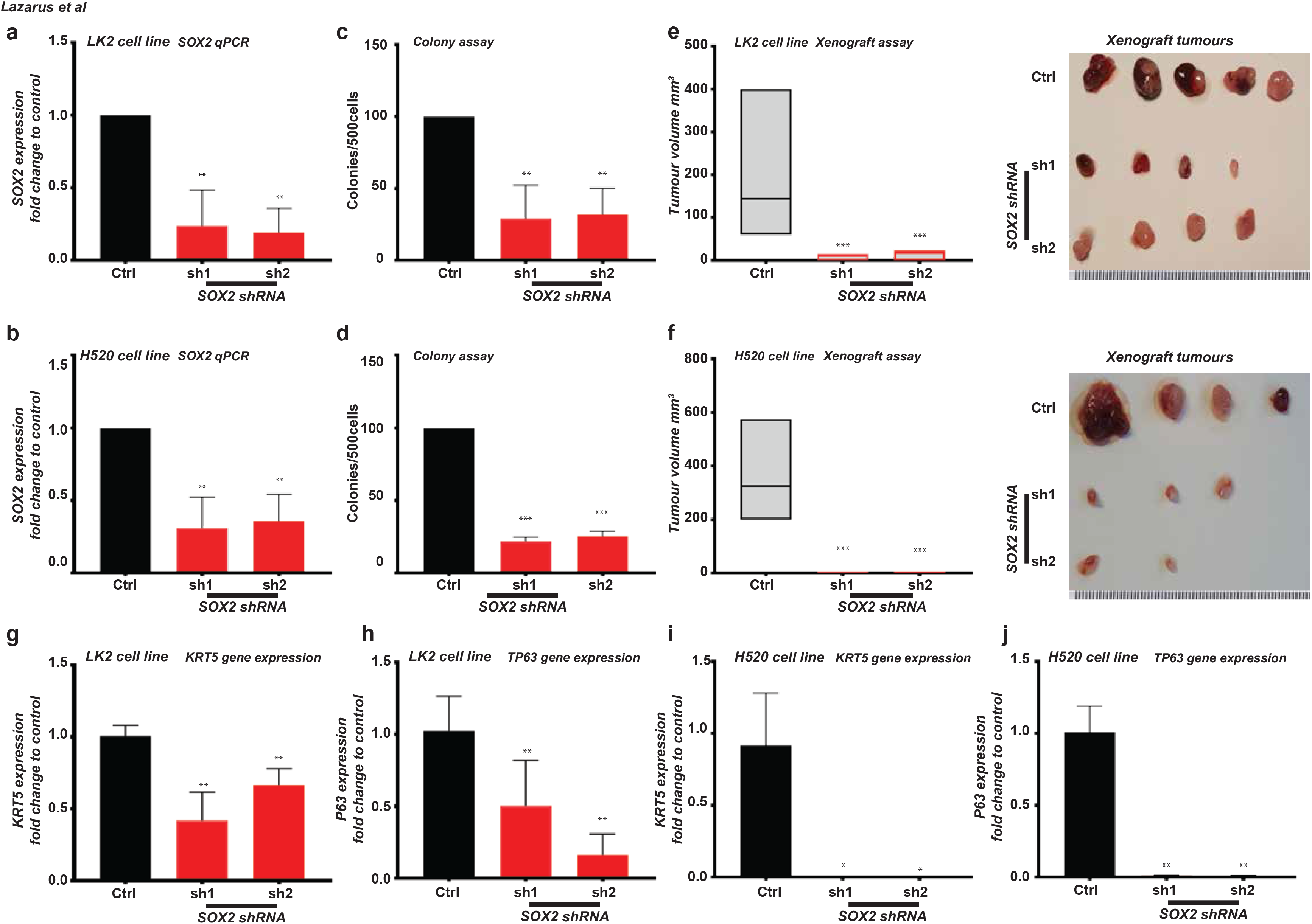
SOX2 KD affects LUSC cell lines tumourigenicity. **(a and b)** Graph showing *SOX2* expression in *SOX2-KD* LK2 (a) and H520 (b) cells. **(c and d)** Graph showing decrease in colony numbers in *SOX2-KD* LK2 **(c)** and H520 **(d)** cells. **(e and f)** Graph depicting reduction in tumour size observed when *SOX2* shRNA1 or shRNA2 transfected LK2 **(e)** and H520 **(f)** cells are injected subcutaneously into mice compared with control. Five (LK2) and four (H520) mice per cell line were monitored for 15 days after which tumours were removed and measured. On the right are images showing actual tumours measured. Data presented as mean ± s.d. One way ANNOVA with post Dunnett test performed, * indicates p<0.05 and ** p<0.005 and *** indicates p<0.001. **(g and h)** *KRT5* expression is reduced in LK2 **(g)** and H520 **(h)** *BCL11A-KD* cells. **(i and j)** *P63* expression is reduced in LK2 **(i)** and H520 **(j)** *BCL11A-KD* cells.

**Supplementary Figure 8.**
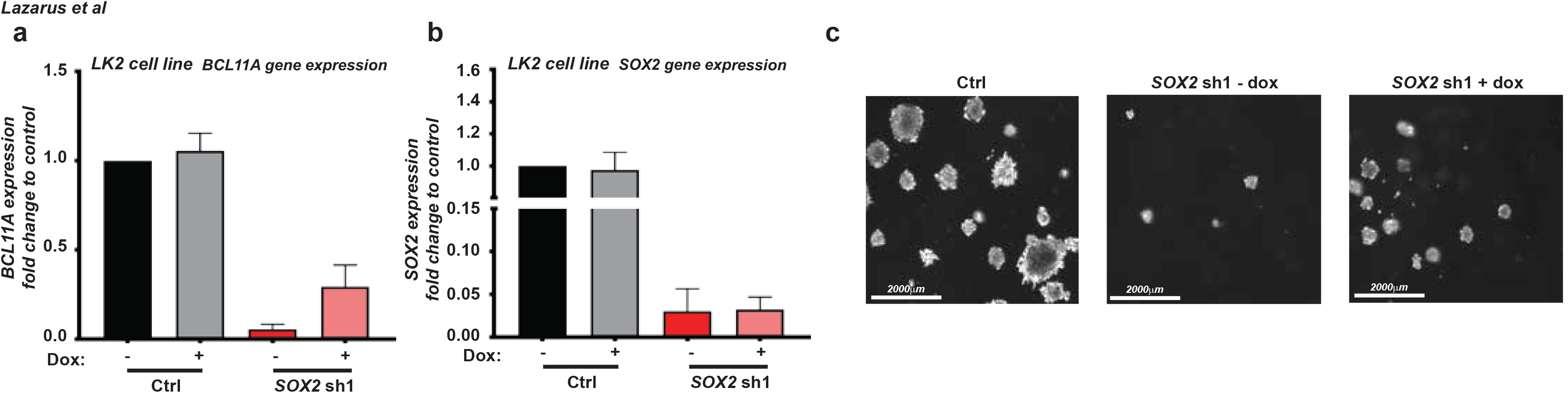
BCL11A is required for SOX2 mediated LUSC phenotype. **(a and b)** Graph showing *BCL11A* **(a)** and *SOX2* **(b)** expression in BCL11A rescue in SOX2-KD cells. Dox inducible *BCL11A* overexpression vector was transfected into control and *SOX2* shRNA1 LK2 cells and Dox treatment was performed for 48 hours. **(c)** Images from 3D matrigel experiment showing reduction in colony numbers after *SOX2-KD* and partial rescue after BCL11A overexpression.

**Supplementary Figure 9.**
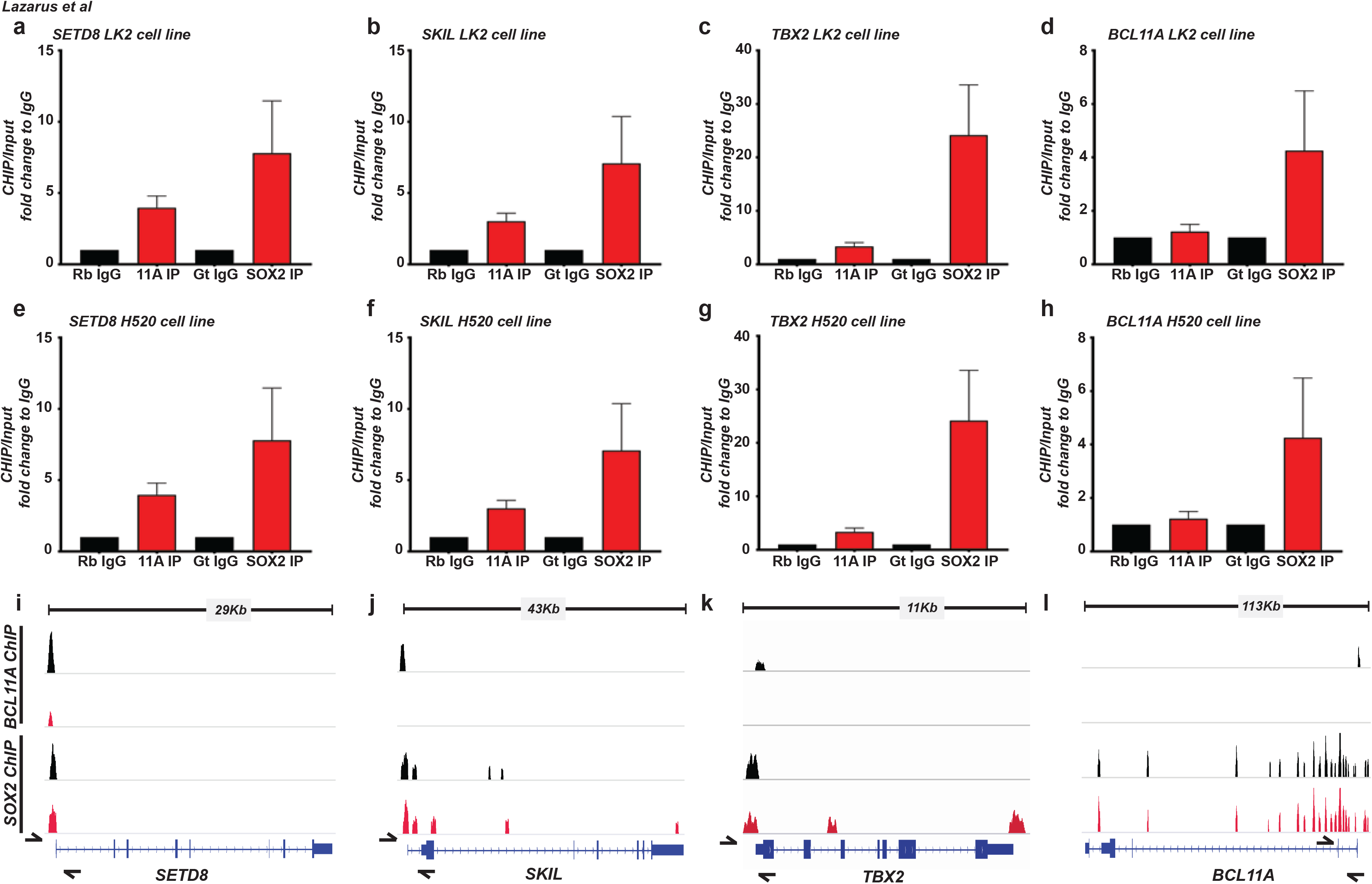
BCL11A and SOX2 bind to common loci on the genome. **(a-h)** ChIP-qPCR showing enrichment for the indicated primers on *SETD8, SKIL, TBX2* and *BCL11A* regions after Chip pull down using anti-BCL11A or anti-SOX2 antibodies in LK2 and H520 cell lines. **(i-l)** Schematic of amplicon locations for ChIP-qPCR validations performed in this study. Arrows represent location of primers used. Error bars represent mean ± s.e.m.; n = 3 (technical replicates).

**Supplementary Figure 10.**
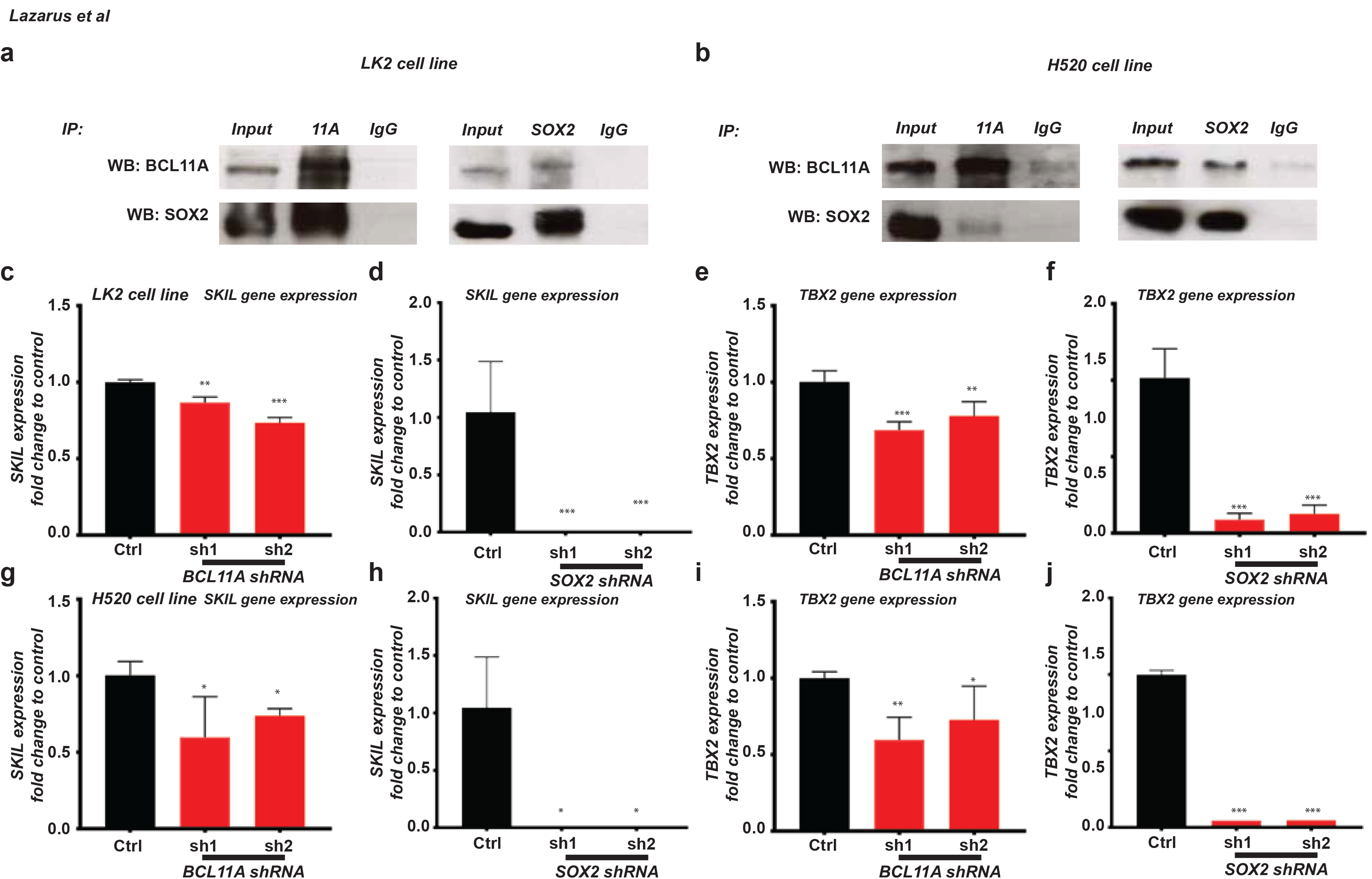
BCL11A and SOX2 co-regulate transcriptional regulators. **(a and b)** Co-Immunoprecipitation of endogenous BCL11A and SOX2 proteins in LK2 **(a)** and H520 **(b)** cell lines. **(c, d, g, h)** *SKIL* gene expression in *BCL11A*-KD or *SOX*-KD LK2 and H520 cell lines. **(e, f, I, j)** *TBX2* gene expression in *BCL11A*-KD or *SOX2*-KD LK2 and H520 cell lines. Data presented as mean ± s.d. (n=3). One way ANOVA with post Dunnett test performed, * indicates p<0.05 and ** p<0.005 and *** indicates p<0.001.

**Supplementary Figure 11.**
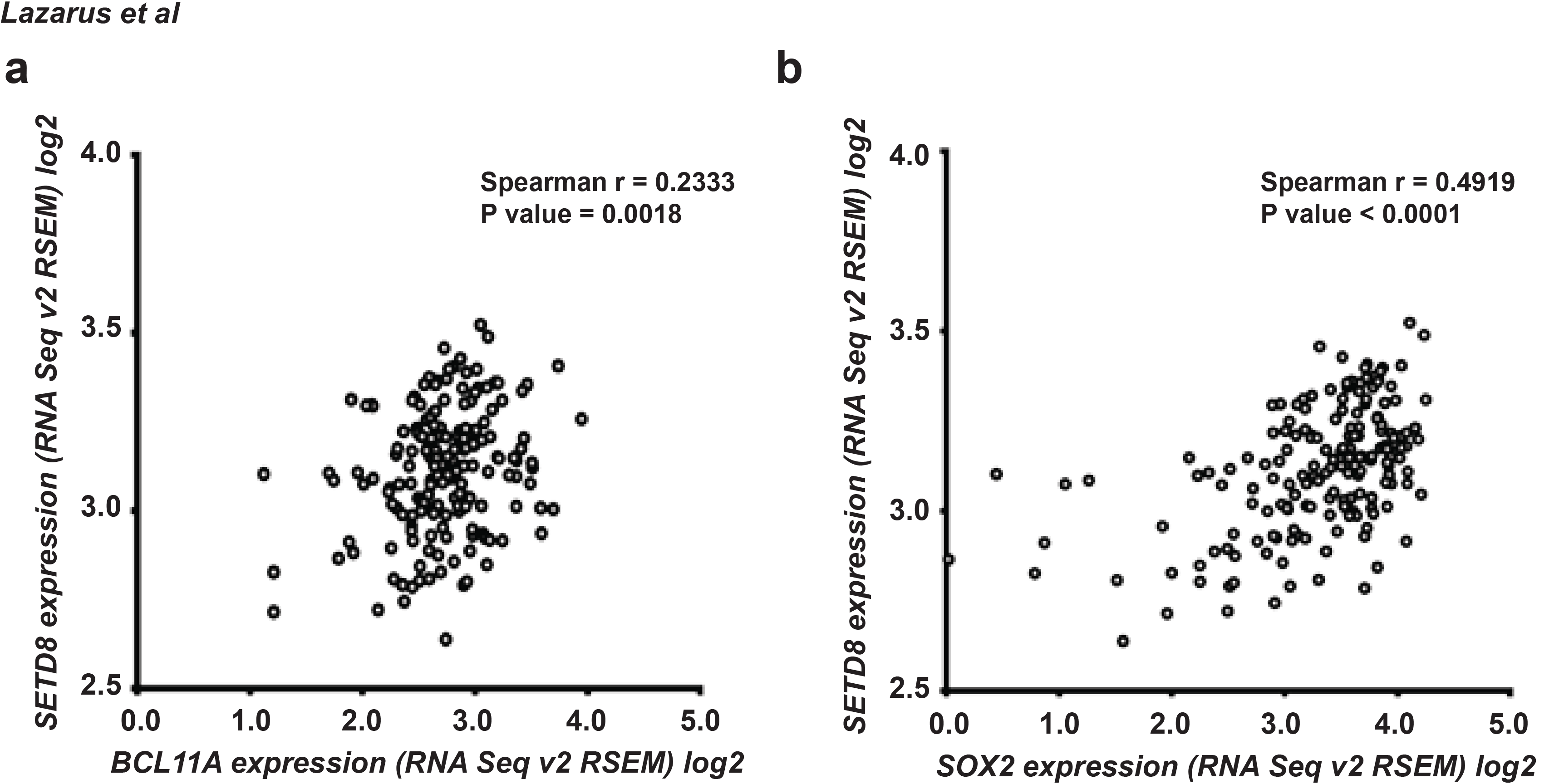
*SETD8* correlates with *BCL11A* and *SOX2* in LUSC patients. **(a)** Scatter plot showing *SETD8* and *BCL11A* expression in TCGA patient tumour samples. **(b)** Scatter plot showing *SETD8* and *SOX2* expression in TCGA patient tumour samples.

## Materials and Methods

### Mouse models

All mice used in this study were maintained at the Sanger Institute or the University of Cambridge. Housing and breeding of mice and experimental procedures were performed according to the UK 1986 Animals Scientific Procedure Act and local institute ethics committee regulations. The *BCL11A^ovx^* allele was generated following a strategy previously described^36^. Briefly, the *ROSA26* allele was targeted with a construct containing human *BCL11A* cDNA preceded by a loxP flanked STOP cassette and marked *eGFP* under the control of an internal ribosomal entry site (IRES) downstream of the inserted cDNA and transgene transcription is controlled by the *CAG* promoter. The generation of the *Bcl11a^cko^* mice was described previously^16^. All mice were 8–12 weeks of age at the time of experiments, and at least 3 mice per cohort were used in each experiment. The primers used for genotyping are listed in Supplementary Table 4.

### Mouse Tracheal Isolation and tracheoshpere culture

Tracheae were incubated in 50 U/ml dispase (Sigma) for 45 minutes at 37°C. 10 ml PBS was injected through each trachea using a 25G 5/8” needle to flush out sheets of epithelial cells. Cells were incubated in 0.25% trypsin for 5 minutes at 37°C. Cells were stained in PBS + 10%FBS + 1:100 anti-EpCAM PE-Cy7 (Biolegend) + 1:5000 DAPI on ice for 20 minutes. Live EpCAM^+ve^ cells were isolated using a MoFlo sorter. 2500 cells were plated in 100μl 1:1 mix of Mouse Tracheal Epithelial Cell (MTEC)/Plus media which is DMEM-Ham’s F-12 (1:1 vol/vol), 15 mM HEPES, 3.6 mM sodium bicarbonate, 4 mMl-glutamine, 100 U/ml penicillin, 100 μg/ml streptomycin, and 0.25 μg/ml fungizone; supplemented with 10 μg/ml insulin, 5 μg/ml transferrin, 0.1 μg/ml cholera toxin, 25 ng/ml epidermal growth factor (Becton-Dickinson, Bedford, MA), 30 μg/ml bovine pituitary extract, 5% FBS, and freshly added 0.01 μM retinoic acid^37^ and growth factor reduced-Matrigel (Corning) per 24-well insert, in duplicate with 500 nM 4-hydroxy tamoxifen (Sigma) or ethanol (vehicle) for each mouse, and cultured for 15 days.

### Cell lines

LK2, NCI-H520, NCI-H157, LUDLU1, SW900, NCI-H1792, NCI-H522, NCI-HCC78, A549, NCI-H1563, NCI-H1975 were all maintained in RPMI 1640 (Gibco), 10% FCS and 1% Penicillin/streptomycin in a 37°C incubator with 5% CO_2_.

### ShRNA mediated knockdown

*BCL11A* shRNA sequences were obtained from TRC consortium (TRCN0000033449 and TRCN0000033453) and cloned into a piggyBac transposon vector (PB-H1-shRNA-GFP) as describe previously^38^. Sox2 shRNA sequences (TRCN0000355694 and TRCN0000257314) were also cloned as above. H520, H1792 and LK2 cells were transfected with 4.0 μg of respective vector using Lipofectamine 3000 or Lipofectamine LTX (Invitrogen). Cells were treated with G418 (400 μg/mL) (Gibco) for 5 days after which GFP^+ve^ cell were sorted using a Sony SH800 cell sorter (Sony, Tokyo, Japan) and cultured.

### Transfection and 3D colony assays

The control or the *BCL11A* overexpression piggybac vectors were delivered into NSCLC cells using Lipofectamine LTX (Invitrogen). Transfected cells were maintained for 48 hours and then cultured in puromycin (1 μg/ml). To induce BCL11A expression in LK2 *SOX2-KD* cells, doxycycline (Clonetech) was used at a final concentration of (1 μg/ml). 3D colony assays were performed by suspending 500 cells in matrigel (BD Biosciences) and seeding this cell-matrigel suspension onto a 6-well plate. The plate was then incubated for 15 minutes in 37°C/5% CO_2_ to allow hardening of suspension. Growth media was added to the well and changed every 2-3 days for 20 days. All experiments were performed in triplicates.

### Preparation of RNA

RNA was extracted using the RNeasy mini kit (Qiagen). Cell cultures in T25 flasks were first washed with cold PBS, and 350 μl of RLT was added. Cells were scraped, passed through a 20G syringe five times and RNA was extracted using the RNeasy mini kit (Qiagen) according to manufacturer instructions. DNA was degraded by adding 20U Rnase-free DnaseI (Roche) for 30 minutes at room temperature. DnaseI treatment was performed on columns.

### Preparation of cDNA and qRT-PCR

1 μg of total RNA was diluted to a final volume of 11 μl. 2 μl of random primers (Promega) were added after which the mixture was incubated at 65°C for 5 mins. A master mix containing Transcriptor Reverse Transcriptase (Roche), Reverse Transcriptase buffer, 2 mM dNTP mix and RNasin Ribonuclease Inhibitors (Promega). This mixture was incubated at 25 °C for 10 minutes, then 42°C for 40 minutes and finally 70°C for 10 minutes. The resulting cDNA was then diluted 1:2.5 in H_2_O for subsequent use. qPCR was performed using a Step-One Plus Real-Time PCR System (Thermofisher Scientific). Either Taqman (ThermoFisher Scientific) probes with GoTaq Real Time qPCR Master Mix (Promega) or primers (Sigma) with PowerUp SYBR Green Master Mix (ThermoFisher Scientific) were used. All probe and primer details can be found in Supplementary Table 5 and 6. The enrichment was normalised with control mRNA levels of GAPDH and relative mRNA levels were calculated using the ΔΔCt method comparing to control group.

### Western Blot

Cells were lysed using RIPA (Cell signalling) and protease inhibitors (Roche) as per manufacturer instructions. Total protein was measured using the bicinchoninic acid (BCA) method (Pierce Biotechnology). In total, 50 mg cell lysate was separated using 7.5% SDS-PAGE gels and transferred to PVDF membranes by electro-blotting. Membranes were blocked in 5% (w/v) milk in Tris-buffered saline containing 0.05% Tween-20 (TBST). Blots were then incubated at 4°C overnight with primary antibodies as indicated, washed in TBST and subsequently probed with secondary antibodies for 1h at room temperature. ECL solution was then added to the membrane and analysed. Antibodies used were, anti-BCL11A (ab191401, Abcam, 1:3000), anti-SOX2 (ab97959, Abcam, 1:2000) and anti-TUBULIN (ab7291, Abcam, 1:10000).

### Co-Immunoprecipitation

Cells were lysed using RIPA (Cell signalling) and protease inhibitors (Roche) as per the manufacturer’s instructions. Total protein was measured using the BCA method as above (Pierce Biotechnology). Briefly, 500μg cell lysates were pre-cleared for 3h at 4°C to remove nonspecific binding. Then, the pre-cleared lysates were incubated with anti-BCL11A (Bethyl, A300-382A) and SOX2 (R&D Systems, AF2018) or control IgG at 4°C overnight. Next day 50μl of Dynabeads Protein G (Thermo Fisher Scientific) were added to each sample. After 3h, the complex was washed three times with RIPA buffer, and then analysed by Western Blot performed as described above.

### Histology, Immunohistochemistry and Immunofluorescence

Cultured organoid were fixed with 4% paraformaldehyde in PBS for 4h at room temperature. After rinsing with PBS, fixed colonies were immobilised with Histogel (Thermo Scientific) for paraffin embedding. 5μm sections of lung tissues or embedded colonies were stained with haematoxylin and eosin (H&E) and immunostained with antibodies for BCL11A (IHC - ab191402, Abcam,1:1000), SOX2 (IHC - ab97959, 1:1000; IF - 14-9811-82, eBioscience, 1:200), Ki67 (IHC - MA5-14520, Thermo Scientific, 1:1000; IF - ab16667, Abcam, 1:300), GFP (IF - ab13970 Abcam, 1:1000), Keratin 8 (IF - TROMA-I, DSHB, 1:100), Keratin 5 (IHC - ab52635, Abcam, 1:1000; IF - 905501, Biolegend, 1:1000), P63 (IHC - ab735, Abcam, 1:200; IF - ab735, Abcam, 1:200), CCP10 (IHC - sc25555, Santa Cruz Biotechnology, 1:500). IHC secondary staining involved an HRP-conjugated donkey anti-rabbit or donkey anti-mouse secondary (1:250, Thermo Scientific) and were detected using DAB reagent (Thermo Scientific). IF secondary staining involved goat anti-chicken 488, goat anti-rabbit 647, goat anti-rat 647, goat anti-rabbit 488 and goat anti-mouse 647 (1:2000, Invitrogen). Nuclear stain was detected using Haematoxylin (IHC) or ProLong Gold Antifade Mountant DAPI (Thermofisher, P36941) (IF). IHC images were acquired using a Zeiss Axioplan 2 microscope and IF images were acquired using a Leica TCS SP5 confocal microscope and analysed on Image J.

### Xenograft tumour assays

One million H520 and LK2 cells were suspended in 25% matrigel (BD Biosciences) and HBSS. This mixture was subcutaneously injected in 6-12 week old NSG mice. To induce BCL11A overexpression in *SOX2-KD* xenografts, mice were fed doxycycline pellets (Envigo, TD.01306, 625mg/kg). Mice were culled as specified in figure legends and resulting tumours were analysed.

### ChIP-Seq and ChIP-qPCR

ChIP-Seq experiments were performed as described^39^. Antibodies used were BCL11A (Bethyl, A300-382A) and SOX2 (R&D Systems, AF2018). Briefly 2 x 15cm plates per cell line were formaldehyde crosslinked, nuclear fraction was isolated and chromatin sonicated using Bioruptor Pico (Diagenode). IP was performed using 100μl of Dynabeads Protein G (Thermo Fisher Scientific) and 10μg of antibody. The samples were then reverse crosslinked and DNA was eluted using Qiagen MinElute column. Sample was then processed either for sequencing or qPCR. Primers used for qPCR are listed in Supplementary Table 7.

### Drug assays

SETD8 inhibitor NSC663284 (Cayman Chemical Company, 13303) was suspended using DMSO in 10mM stock concentration. Cisplatin (LKT Laboratories, C3374) was suspended in 154mM NaCl at a 3mM stock concentration. Cells were cultured as above and seeded at 1000 cells per 96-well plate and left to recover for 24h. The edges of the 96-well plate were avoided to ensure accuracy in measurement. For NSC663284, an initial dilution of 1:100 from stock was performed in RPMI media for the first concentration of 10-4M. Half log dilutions were performed in RPMI media reaching 10-6M and after which full log dilutions were performed reaching 10-10M. For cisplatin, initial dilution in RPMI media were made to achieve 100μM and 75μM. The 100μM solution was used to make the following solutions 50μM, 25μM, 12.5μM, 6.75μM, 3.75μM, 1.5μM and 0.5μM. Cells were treated with vehicle for 48h after which the above doses of cisplatin were added. For cisplatin + NSC663284 experiment the NSC663284 IC50 concentration for each cell line was calculated and added initially for 48hrs and then added again along with the cisplatin dilution series as above for 24hrs. Data analysis for drug inhibitor assays was performed in GraphPad Prism 7.02 (San Diego, CA). Data were fitted to obtain concentration-response curves using the three-parameter logistic equation (for pIC50 values). Emax was constrained to 100% while the basal (Emin) parameter was contrained to 20%. Statistical differences were analyzed using one-way ANOVA or Student's t test as appropriate with post hoc Dunnett’s multiple comparisons, and p < 0.05 was considered significant.

### BCL11A IHC on patient tumours

TMAs contained LUAD (n=99) and LUSC (n=120) cases of archival primary pulmonary tumours collected under East Midlands NRES REC approved project (ref. 14/EM/1159). 1mm (n=3) cores are present per case which were initially scored (average nuclear staining intensity) as 0=neg, 1=weak, 2=moderate, 3=strong. A median score was calculated for each case and re classified as 0=neg, 1=moderate, 2=strong. All tissues and data are anonymised to the research team. IHC was performed using BCL11A antibody (ab19487, 1:200) with CC1 antigen retrieval using a Ventana discovery xt. Digital images of stained TMAs were scanned with a Nanozoomer RX instrument and scored on-screen by MD.

### TCGA gene expression analysis

The TCGA data was accessed from the recount2 database^40^ containing gene level count data from RNA-seq and clinical data from primary tumor samples from patients diagnosed with LUAD and LUSC, respectively. EdgeR^41^ was used to test for differential expression of transcription factors (as defined by Tfcheckpoint.org). For this, compositional differences between samples were normalized using the trimmed median of M-values method and a gene-specific dispersion was estimated for each gene. A negative binomial generalized log-linear model was fit to each gene with the covariates “plate+disease_type” (for the LUAD versus LUSC comparison) or “patient+disease_type” (for the tumor versus normal comparison). A log-likelihood ratio test was conducted to test whether the coefficient of the disease_type variable is non-zero, followed by Benjamini-Hochberg adjustment of P values to account for multiple testing. Plate A277 and A278 from the LUAD dataset were removed prior to analysis as they showed a clear separation along the first component in a principal component analysis from all other LUAD samples.

### ChIP-Seq analysis

ChIP-Seq libraries were sequenced on illumina Hiseq2000 platform at the Wellcome Sanger Institute. Each library was divided into two and sequenced on different lanes. Reads were subsequently run through a pipeline at the sequencing facility to remove adapter sequences and align to the reference genome among others. Alignment was done using mem algorithm in BWA (version 0.7.15) and human_g1k_v37 was used as the reference genome. Aligned and processed reads were received as compressed CRAM files. Samtools (version 1.3.1) was used to decompress the CRAM files and filter uniquely mapped reads in proper pairs. Next, reads from the two runs were combined into a single BAM file using ‘merge’ function in samtools. Bedtools intersect was then used to remove reads falling into blacklisted genomic regions or unplaced genomic contigs of the GRCh37 assembly before marking and removing duplicate reads using MarkDuplicates function in Picard tools. Next, DownsampleSam in picard tools was used to sample ~105 million reads from each BAM file. Significantly enriched genomic regions relative to input DNA were identified using MACS2 (version 2.1.1.20160309) with p-value cutoff of 1.00e-05. Heatmaps generation: Mapped read counts were calculated in a 10bp window and normalised as RPKM (Reads Per Kilobase per Million mapped reads) using bamCoverage module from deeptools (version 2.5.1)^42^. This coverage file was used to compute score matrix ±1kb around peak summits using computeMatrix reference-point module (from deeptools version 2.5.1)^42^. Heatmaps of binding profiles around peak summits were then generated using plotHeatmap module in deeptools (version 2.5.1)^42^. Number of overlapping peaks between BCL11A and SOX2 and nearest downstream genes to peaks were determined using ODS and NDG utilities respectively in PeakAnnotator (version 1.4). For annotating nearest downstream genes, Homo sapiens GRCh37 (release 64) from ensembl was used.

## Author contribution

K.L. designed and performed the majority of the experiments and analysed most of the data. F.H. analysed the ChIP-seq data. E.Z. performed the ChIP-qPCR experiments. K.B. analysed the TCGA data. S.P. performed some cell line work. R.U assisted with the BCL11A rescue experiment. M.F.S performed Co-IP experiment. L.B. measured the airway thickness. L.S.C. characterised the lung pathology of adenovirus mice. G.L designed and analysed the drug assays. J.K and J.H.L performed the tracheoshpere organoid experiment. L.L.C and F.M performed the mouse tissue IHC. M.D and J.LQ performed and analysed the BCL11A IHC on patient tumours. P.L. assisted with the NGS sequencing and provided the *Bcl11a^cko^* mice. Adenovirus Cre administration was performed under G.E. supervision. D.C. generated the *BCL11A^ovx^* mice. W.T.K conceptualised and supervised the study. K.L and W.T.K wrote the manuscript.

## Acknowledgements

We would like to thank the staff at Sanger Institute, Research Service Facility (RSF) for their assistance. We would like to thank Dr. Catherine Wilson and Dr. Deborah Burkhart (Department of Biochemistry, Cambridge) for her help with the Adenovirus experiment. We would like to thank Dr. Emma Rawlins for helpful discussions and comments. M.F.S was funded by Associazione Italiana per la Ricerca sul Cancro (AIRC, 16719/2015). W.T.K is funded by a CRUK career establishment award (C47525/A17348), University of Cambridge and Magdalene College, Cambridge. We would also like to thank the Isaac Newton Trust for funding. G.L. was supported by the BBSRC (Grant BB/M00015X/2).

